# The TRF2 General Transcription Factor Is a Key Regulator of Cell Cycle Progression

**DOI:** 10.1101/2020.03.27.011288

**Authors:** Adi Kedmi, Anna Sloutskin, Natalie Epstein, Lital Gasri-Plotnitsky, Debby Ickowicz, Irit Shoval, Tirza Doniger, Eliezer Darmon, Diana Ideses, Ziv Porat, Orly Yaron, Tamar Juven-Gershon

**Affiliations:** The Mina and Everard Goodman Faculty of Life Sciences, Bar-Ilan University, Ramat Gan 5290002, Israel; The Flow Cytometry Unit, Life Sciences Core Facilities, Weizmann Institute of Science, Rehovot 7610001, Israel

**Keywords:** Basal transcription machinery, RNA polymerase II, TATA box-binding protein (TBP), TBP-related factor 2 (TRF2), cyclin genes

## Abstract

TRF2 (TATA-box-binding protein-related factor 2) is an evolutionarily conserved general transcription factor that is essential for embryonic development of *Drosophila melanogaster, C. elegans*, zebrafish and *Xenopus*. Nevertheless, the cellular processes that are regulated by TRF2 are largely underexplored.

Here, using *Drosophila* Schneider cells as a model, we discovered that TRF2 regulates cell cycle progression. Using flow cytometry, high-throughput microscopy and advanced imaging-flow cytometry, we demonstrate that TRF2 knockdown regulates cell cycle progression and exerts distinct effects on G1 and specific mitotic phases. RNA-seq analysis revealed that TRF2 regulates the expression of *Cyclin E* and the mitotic cyclins, *Cyclin A, Cyclin B* and *Cyclin B3*, but not *Cyclin D* or *Cyclin C*. To identify proteins that could account for the observed regulation of these cyclin genes, we searched for TRF2-interacting proteins. Interestingly, mass spectrometry analysis of TRF2-containing complexes identified GFZF, a nuclear glutathione S-transferase implicated in cell cycle regulation, and Motif 1 binding protein (M1BP). Furthermore, available ChIP-exo data revealed that TRF2, GFZF and M1BP co-occupy the promoters of TRF2-regulated genes. Using RNAi to knockdown the expression of either M1BP, GFZF, TRF2 or their combinations, we demonstrate that although GFZF and M1BP interact with TRF2, it is TRF2, rather than GFZF or M1BP, that is the main factor regulating the expression of *Cyclin E* and the mitotic cyclins. Taken together, our findings uncover a critical and unanticipated role of a general transcription factor as a key regulator of cell cycle.

## Introduction

Multiple biological processes and transcriptional programs are regulated by RNA polymerase II (Pol II). The initiation of transcription of protein-coding genes and distinct non-coding RNAs occurs following the recruitment of Pol II to the core promoter region by the general/basal transcription machinery [1–4]. The core promoter, which directs accurate initiation of transcription and encompasses the transcription start site (TSS), may contain short DNA sequence elements/motifs, which confer specific properties to the core promoter [1, 4–10]. The first step in the recruitment of Pol II to initiate transcription is the binding of TFIID, which is composed of TATA-box-binding protein (TBP) and TBP-associated factors. Remarkably, although TBP is considered a universal general transcription factor, robust Pol II transcription is observed in mouse TBP-/- blastocysts, indicating the existence of TBP-independent Pol II transcription *in vivo* [11]. The complexity of transcription is also manifested by the existence of diverse transcriptional regulators, among which are the TBP family members. There are three TBP family members in *Drosophila melanogaster:* TBP, TRF1 and TRF2 (reviewed in [12–16]). TRF1, the first *Drosophila* TBP family member identified, is insect specific [17]. An evolutionary conservation analysis indicated that TRF2 (also known as TLP (TATA-like protein), TLF (TBP-like factor), TRP (TBP-related protein) and TBPL1 (TBP-like 1)), is highly conserved in evolution [12, 18–22] and is present in all bilaterian organisms, but not in any of the non-bilaterian genomes available [19]. It was further discovered that TRF2, which is involved in Pol II transcription, evolved by duplication of the TBP gene [19]. Yet, unlike TBP and TRF1, TRF2 does not bind TATA-box containing promoters [19, 20, 22]. There are two *Drosophila* TRF2 protein isoforms that result from an internal translation initiation: the evolutionarily conserved short isoform (632 aa; typically referred to as “TRF2”) and a long *Drosophila*-only isoform (1715 aa), in which the same short amino acid sequence is preceded by an N-terminal domain [23]. TRF2 affects early embryonic development, differentiation and morphogenesis of *Drosophila, C. elegans*, zebrafish and *Xenopus* [23–32]. Mouse TRF2 is essential for spermiogenesis [33–35].

One of the open questions in the transcriptional regulation field is what are the cellular functions of TRF2. Despite its importance in development, the cellular processes that are regulated by TRF2 remain largely underexplored. To identify and characterize the cellular processes that are regulated by TRF2, we used *Drosophila* S2R+ cells as a model and knocked down the expression of TRF2. We discovered that reduced expression of TRF2 (but not TBP) exerts distinct effects on G1, G2/M and specific mitotic phases, as demonstrated by quantitative high-throughput imaging flow cytometry. We further discovered that TRF2 controls the expression of *Cyc E* and the mitotic *Cyc A, Cyc B* and *Cyc B3* genes. Using mass spectrometry analyses of TRF2-interacting proteins and available ChIP-exo data, we demonstrate the co-occupancy of TRF2, GFZF (GST-containing FLYWCH zinc-finger protein) and M1BP (motif 1 binding protein) in the majority of promoters bound by each of the three factors. Remarkably, the promoters of the TRF2-regulated mitotic cyclins and *Cyclin E* are bound by the three factors, whereas the promoters of *Cyclin C* and *Cyclin D*, which are not regulated by TRF2, are not bound. Furthermore, the Motif 1 sequence element is enriched in the promoters of genes bound by all three proteins, suggesting the involvement of GFZF and M1BP as co-factors in TRF2-regulated cell cycle progression. Moreover, we demonstrate that TRF2, rather than M1BP or GFZF, is the main factor that regulates the expression of the mitotic cyclins and *Cyclin E*. Importantly, while general/basal transcription factors might be viewed as having a somewhat “generic” role, our findings emphasize the unique, unanticipated functions of *Drosophila* TRF2 as an essential factor for cell cycle progression and apoptotic cell death.

## Results

### Knockdown of endogenous TRF2 expression results in altered cell cycle distribution and G1 arrest

The TBP-related transcription factor TRF2 is a key general/basal transcription factor (reviewed in [12–16]), yet the cellular processes that are regulated by TRF2 remain largely underexplored. To investigate the cellular functions of TRF2, we used *Drosophila* S2R+ cells as a model, and knocked down its expression by RNAi using non-overlapping dsRNA probes (Fig. S1a). TBP knockdown was used as a control throughout this study. The resulting reduction in protein expression was verified by western blot analysis (Fig. S1b, c).

To identify the targets that are unique to TRF2, we used RNAi to knockdown either TRF2 or TBP in *Drosophila* S2R+ cells, and performed RNA-seq analysis at two time points, 48h and 72h. We then performed Gene Ontology (GO) terms analysis of genes that were either downregulated or upregulated following TRF2 knockdown (using string.db.org) in order to identify specific cellular processes that are regulated by TRF2. Surprisingly, we discovered that genes that were downregulated following TRF2 knockdown are enriched for cell cycle and mitotic cell cycle processes, while genes that were upregulated following TRF2 knockdown are enriched for response to stimulus and stress (Table S1). Interestingly, while the number of genes that were downregulated was only half of the number of the upregulated genes following TRF2 knockdown (337 vs. 684 genes, respectively), the enrichment scores (−log_10_(P values)) of the downregulated genes were 5-fold higher. We thus decided to examine the effects of TRF2 knockdown on cell cycle distribution. To knockdown the expression of the endogenous genes, S2R+ cells were incubated for three days with dsRNA probes against *Trf2, Tbp* or *exd*, as a negative control. Cells were harvested, fixed and analyzed by flow cytometry. Control S2R+ cells display a normal profile with an average of ~31% of cells in G1 phase and with an average of ~34% of cells in G2/M phase, similarly to mock treated cells (which were processed similarly, but were not incubated with any dsRNA; Fig. 1). Following TRF2 knockdown by either one of the four dsRNA probes, we observed a distinct decrease in the fraction of cells in G2/M phase (an average of ~20%), as well as the fraction of cells in S phase (an average of ~20%), with a concomitant increase in the fraction of cells in G1 phase (an average of ~55%). Remarkably, these effects are unique to TRF2 knockdown, as the knockdown of TBP resembles the cell cycle distribution of control and mock treated cells (Fig. 1).

**Fig. 1.**
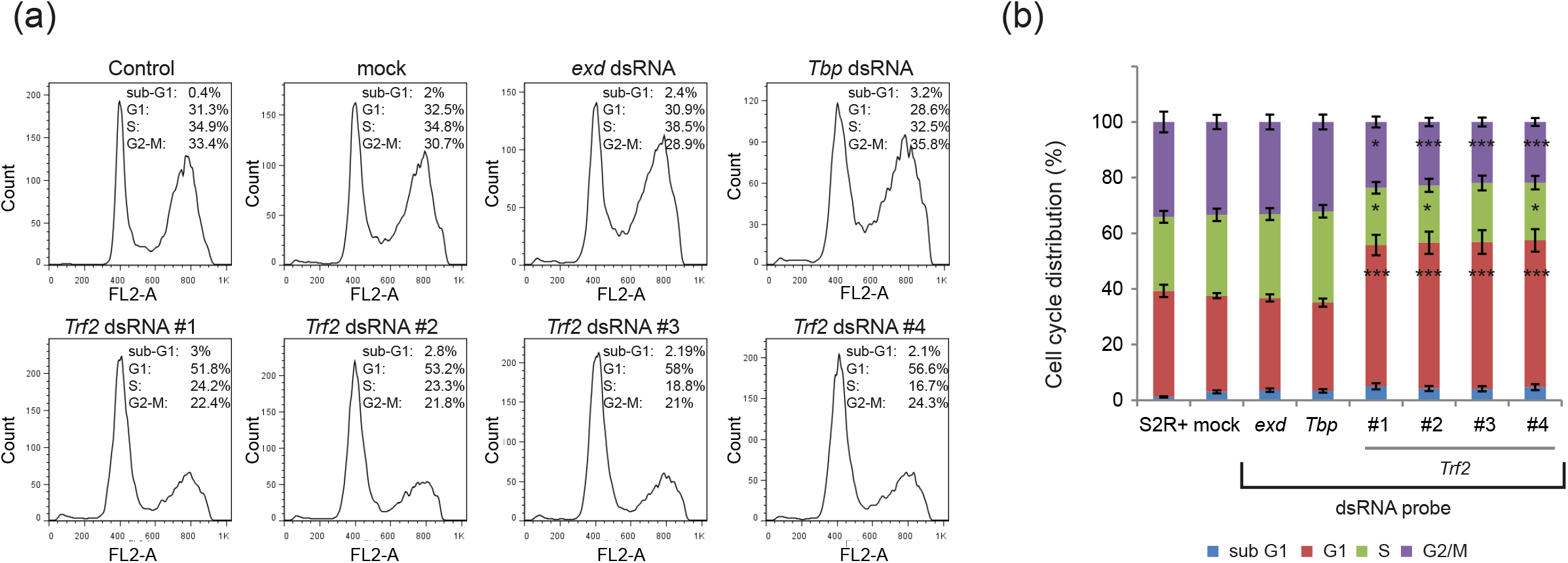
Knockdown of *Trf2*, but not *Tbp*, affects cell cycle distribution. *Drosophila* S2R+ cells were incubated for three days with dsRNA directed against *Trf2, Tbp* and *exd*. Cells were fixed with 80% ethanol and stained with Propidium-Iodide (PI) for flow cytometry (FACS) analysis. (a) Cell cycle distribution histograms of a representative experiment. (b) Average cell cycle distribution determined by FACS analyses of eight independent experiments (*0.01 < *p* ≤ 0.05, **0.005 ≤ *p* ≤ 0.01, \*\*\**p* < 0.005, two-tailed Students t-test; comparison to mock).

Our RNA-seq analysis reveals subsets of genes that are involved in S and G2/M phases. In addition, the cell cycle analysis indicates that TRF2 plays a role in S and/or G2/M phases. To determine if TRF2 affects cell cycle progression to S phase, S2R+ cells were arrested in G1 with 1mM hydroxyurea (HU) for 18h following knockdown of either TRF2 (dsRNA probes #1 and #2) or TBP, and then released to cycle by replacing the medium with fresh medium. As can be seen in Fig. 2, HU treatment (0h) resulted in accumulation of cells in G1 (~55%). Two hours following the release, mock treated cells returned to cycle (indicated by the decrease in the number of cells in G1), and the fraction of cells in S and G2/M increased. Similarly, cells in which TBP was depleted by RNAi, returned to cycle. Surprisingly, unlike mock or TBP RNAi-treated cells, cells in which TRF2 was depleted by either one of the two probes, remained in G1 (~55%) and did not return to cycle (Fig. 2, Fig. S2). Moreover, even 8h following the removal of HU, cells in which TRF2 was knocked-down remained G1-arrested. Our findings imply that endogenous TRF2 is involved in progression into S phase.

**Fig. 2.**
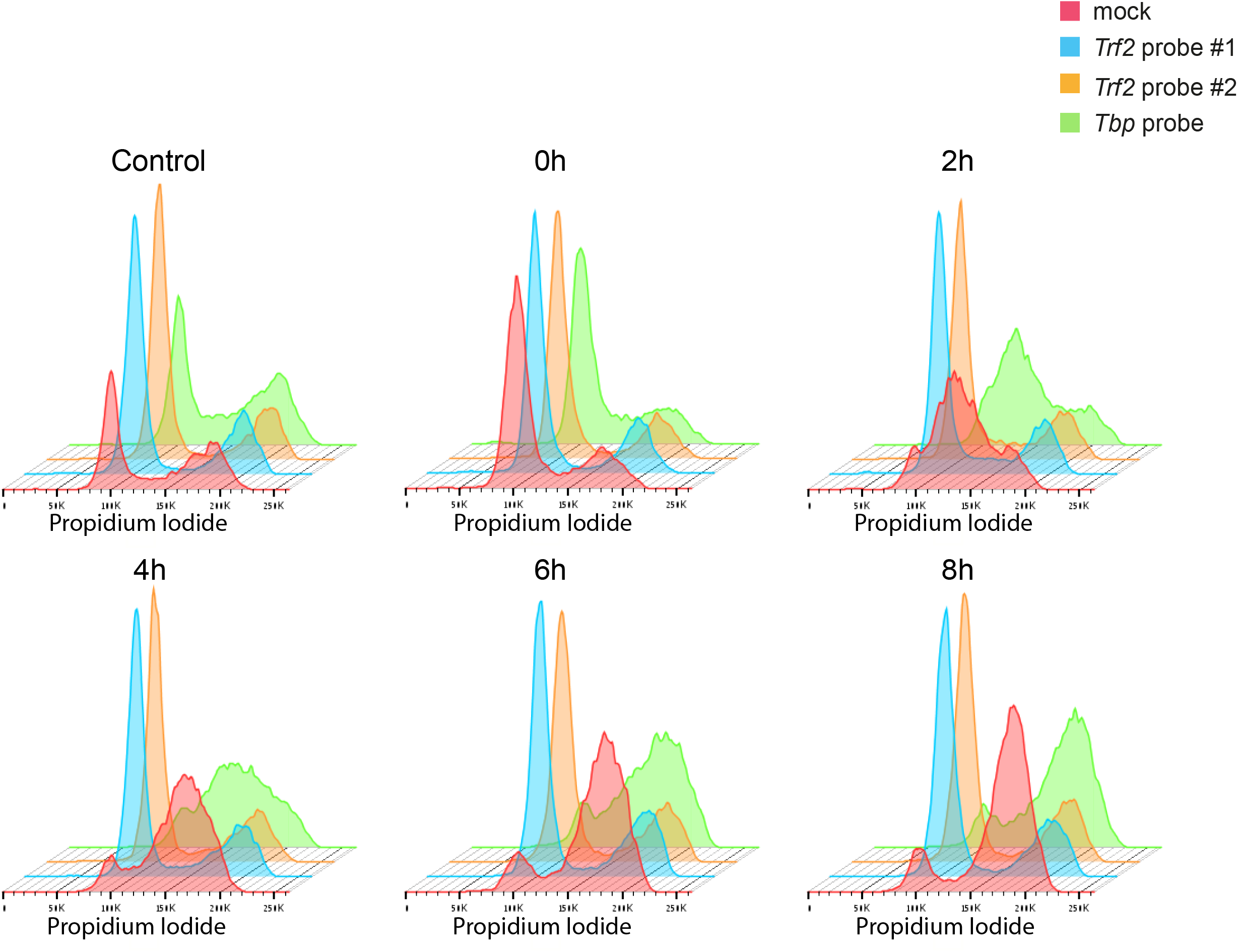
TRF2 is involved in S phase progression. *Drosophila* S2R+ cells were incubated for three days with dsRNA probes directed against *Trf2* or *Tbp*. Next, cells were either left untreated or treated with 1mM Hydroxyurea for 18h. The cells were allowed to resume cell cycle for 2h, 4h, 6h or 8h in fresh medium containing 40μM 5-Bromo-2’-deoxyuridine (BrdU), and were then fixed with 80% ethanol overnight and analyzed by FACS using BrdU-PI staining. Each histogram plots the PI fluorescence intensity (representing DNA content) on the X-axis, and cell count on the Y-axis.

### TRF2 regulates the expression of specific cyclin genes

To further explore the connection between TRF2 and cell cycle progression, we turned to our RNA-seq data for cell cycle-related genes that may be influenced by knockdown of TRF2. We discovered that following TRF2 knockdown, the expression of the *Cyc A, Cyc B, Cyc B3* and *Cyc E* cell cycle regulators was significantly reduced (over 2.3-fold). Notably, the expression levels of *Cyc C* and *Cyc D* were unchanged (Table S1). In order to verify the effect of TRF2 knockdown on the expression of these genes, reverse transcription-qPCR analysis of endogenous cyclin genes was performed on mock, TRF2 (probes #1 and #2), exd or TBP RNAi-treated cells. Knockdown of TRF2 by either probe #1 or #2 significantly reduced the expression of *Cyc A, Cyc B* and Cyc *B3*, as compared to mock treated cells, much more than knockdown of TBP (Fig. 3a). As there was a difference between the effects of probe #1 and #2 on *Cyc E* expression, 4 non-overlapping TRF2 dsRNA probes were used to assess the effect of TRF2 knockdown on *Cyc E* expression (Fig. 3b). Notably, each of the 4 non-overlapping TRF2 dsRNA probes reduces *Cyc E* expression. Unlike *Cyc D* (which is required for G1 progression [36, 37]), the expression of *Cyc E* (which promotes G1-S transition [38, 39]) was reduced following TRF2 knockdown (Fig. 3a, b). We next tested whether TRF2 knockdown affects the protein levels of Cyc A, Cyc B and Cyc E. Unfortunately, we were unable to detect endogenous Cyc A and Cyc B protein expression using publicly available anti-*Drosophila* Cyc A and Cyc B antibodies (data not shown). Remarkably, using anti-*Drosophila* Cyc E antibodies, we observed a distinct reduction in Cyc E protein levels following knockdown of TRF2 (but not TBP), using both TRF2 probes (Fig. 3c), further suggesting that TRF2 regulates G1-S transition by modulating the expression of *Cyc E*.

**Fig. 3.**
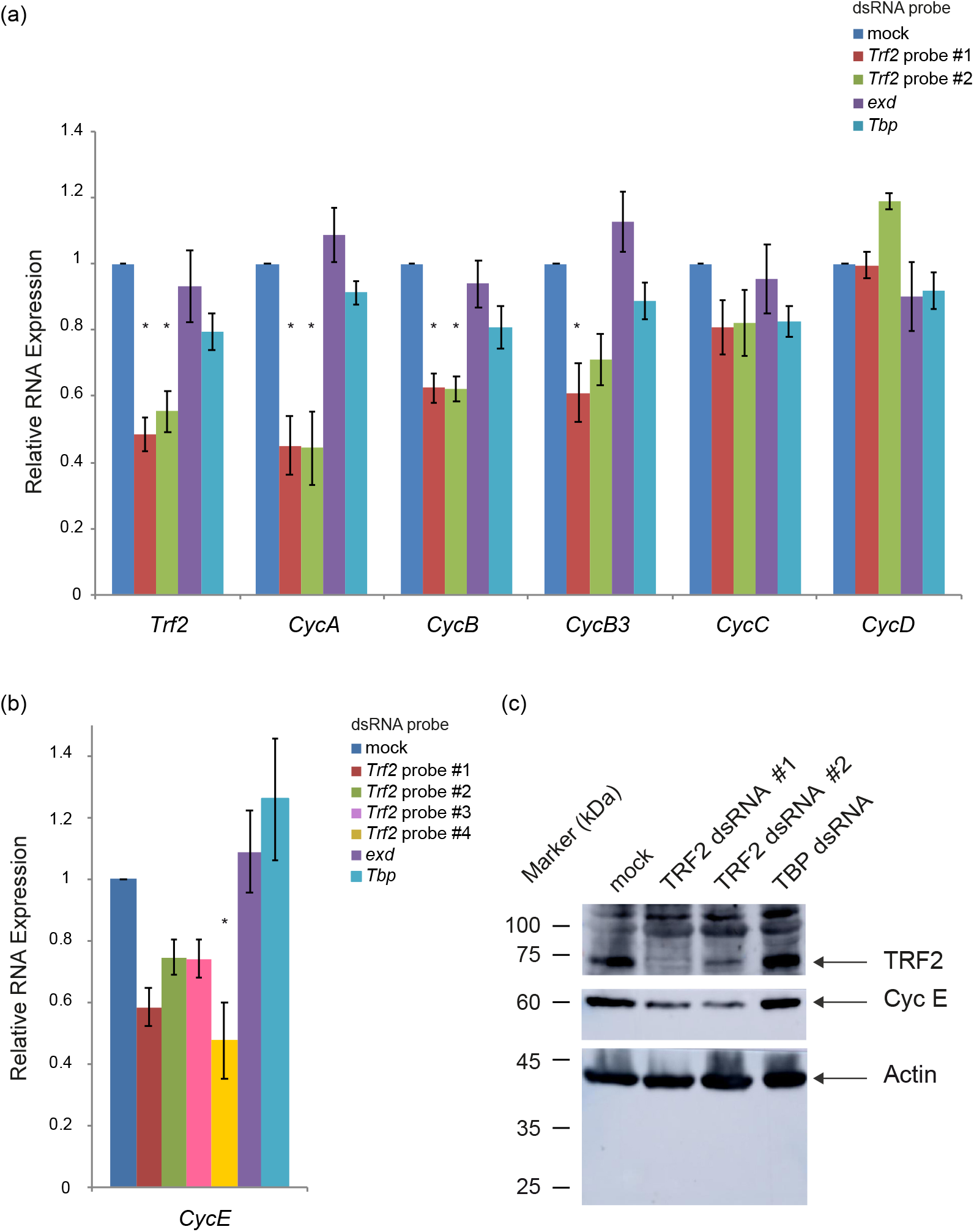
Knockdown of *Trf2* expression by RNAi reduces the expression of specific cyclin genes. *Drosophila* S2R+ cells were incubated for three days with dsRNA probes directed against *Trf2, exd* and *Tbp*. RNA was isolated from the cells and reverse transcribed (RT) to cDNA. Real-time PCR (qPCR) experiments were used to analyze the RNA levels of the endogenous genes: (a) *Trf2, CycA, CycB, CycB3, CycC, CycD* and (b) *CycE*. As there were differences in *CycE* expression following RNAi with probe #1 compared to probe #2, four non-overlapping probes were used to knockdown the levels of *Trf2* expression towards the analysis of *CycE* expression. qPCR experiments were performed in triplicates, and the graph represents the average of three to eight experiments. Error bars represent the SEM. *p< 0.05, oneway ANOVA followed by Tukey’s post hoc test as compared to the mock treatment of the relevant gene. (c) Western blot analysis following TRF2 and TBP knockdown in S2R+ cells, using anti-TRF2 polyclonal antibodies and anti-Cyc E monoclonal antibodies. Actin was used as a loading control.

### TRF2 regulates distinct mitotic phases

We were intrigued by the downregulation of the mitotic cyclin gene expression (*Cyc A*, *Cyc B* and *Cyc B3*) following TRF2 knockdown (Fig. 3a) and decided to explore the effect of TRF2 downregulation on mitotic phases. To this end, we knocked-down the expression of TRF2 or TBP and stained cells for mitotic chromatin (anti-phospho-Histone H3 (Ser10)), DNA (Hoechst) and filamentous Actin (Phalloidin). These allowed us to analyze the fraction of cells in mitosis following TRF2 knockdown (Fig. 4a). To determine the number of mitotic cells, we developed a pipeline to automatically detect the Hoechst and phospho-Histone H3 signals. Notably, there was a reduction in the number of cells undergoing mitosis following TRF2 knockdown, as compared to mock or TBP RNAi treated cells (Fig. 4b).

**Fig. 4.**
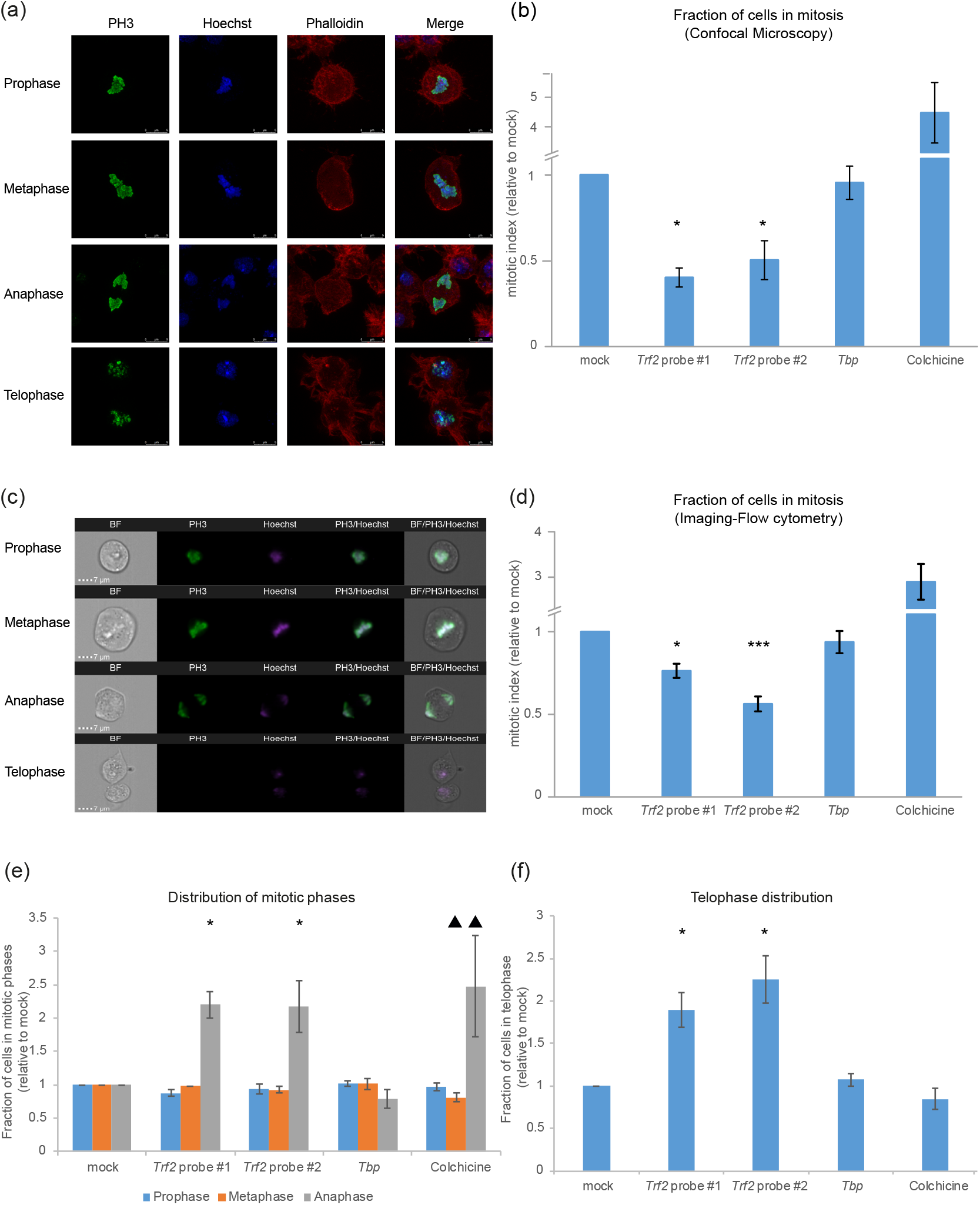
TRF2 regulates cell cycle progression through mitosis. *Drosophila* S2R+ cells were incubated for three days with dsRNA directed against TRF2 or TBP. Cells were then fixed with 4% PFA and stained with a phospho-Histone H3 (Ser10) antibody (PH3, mitotic marker; green), Hoechst (DNA visualization; Blue) and Phalloidin (filamentous Actin visualization; red). (a) Representative confocal microscopy images of *Drosophila* S2R+ cells in different mitotic phases. (b) Comparison of mitotic indices following each treatment, calculated based on microscopic analysis. Shown are the averages of three independent experiments, in which a total of 100,000-800,000 cells were analyzed for each treatment (*0.01 < *p* ≤ 0.05, **0.005 ≤ *p* ≤ 0.01, \*\*\**p* < 0.005, two-tailed Students t-test; comparison to mock). (c) Representative images obtained by imaging flow cytometry analysis. (d) Comparison of mitotic indices following each treatment, calculated based on imaging flow cytometry. Shown are the averages of three independent experiments, in which a total of 40,000 cells were analyzed for each treatment (*0.01 << *p* ≤ 0.05, **0.005 ≤ *p* ≤ 0.01, \*\*\**p* < 0.005, two-tailed Students t-test; comparison to mock). (e-f) Distribution of mitotic phases among all mitotic cells, based on imaging flow cytometry. Shown are the averages of three independent experiments, in which a total of 40,000 cells were analyzed for each treatment (*0.01 < *p* ≤ 0.05, **0.005 ≤ *p* ≤ 0.01, \*\*\**p* < 0.005, two-tailed Students t-test; comparison to mock). Filled triangles indicate aberrant chromosomal morphology in Colchicine-treated cells. PhosphoHistone H3 Ser10-positive cells (e), were analyzed separately from the PhosphoHistone H3 Ser10-negative cells undergoing mitosis (f). Phospho-Histone H3 Ser10negative cells undergoing mitosis (as identified by the imaging flow cytometer) were defined as cells in telophase (f). Notably, cell counts of telophase cells likely include Phospho-Histone H3 Ser10-negative doublet cells, which, even using the high-resolution Imagestream, could not be distinguished from telophase cells.

To explore the effect of TRF2 on G2/M, we sought to synchronize cells in G2/M. Unfortunately, we were unable to synchronize cells in G2/M in a reversible manner (see methods), and thus we could not perform G2/M block-release experiments. Nevertheless, we succeeded in discerning the effects of TRF2 on specific mitotic phases by employing advanced imaging-flow cytometry analysis (ImageStreamX mark II imaging flow-cytometer, Amnis Corp, Seattle, WA, Part of EMD Millipore). Imaging-flow cytometry analysis combines the high-quality imaging and functional insights of microscopy with the speed, sensitivity, and phenotyping abilities of flow cytometry. We knocked down the expression of TRF2 or TBP and stained cells with anti-phospho-Histone H3 (Ser10) antibodies and Hoechst. A total of 40,000 cells of each treatment were analyzed by an ImageStream flow cytometer to determine the number of cells in each mitotic phase, according to their nuclear morphology (Fig. 4c-f, Fig. S3, Table S2). Remarkably, although 40,000 cells were analyzed in each experiment, only a few hundred cells were mitotic, and following TRF2 knockdown there was an even bigger reduction in the total number of mitotic cells (Fig. 4b, d). Notably, despite the overall reduction in the mitotic cell population, knockdown of TRF2 (but not TBP) resulted in a significant accumulation of cells in anaphase and telophase (Fig. 4e, f).

To validate the accurate identification of mitotic cells, we used Colchicine as a control. Colchicine treatment resulted in accumulation of cells in mitosis, specifically in anaphase (Fig. 4e). Multiple studies have shown that Colchicine disrupts the metaphase to anaphase transition (see for example, [40]), yet, the increase in anaphase has also been documented [41]. Furthermore, it is established that different cell types behave differently during mitosis in the presence of drugs that disrupt microtubules function [42]. Interestingly, morphological examination of the Colchicine-treated S2R+ cells indicated aberrant DNA staining patterns and mal-oriented clumped chromosomes in prophase, metaphase and anaphase cells (Fig. S4), in line with the absence of a spindle. Notably, mal-oriented un-centered chromosomes, such as those observed in Colchicine-treated cells, were not observed in TRF2-RNAi treated cells.

### The effects of TRF2 knockdown on cyclin gene expression correlates with the promoter occupancies of TRF2 and its co-factors GFZF and M1BP

To better understand how TRF2 regulates the expression of the cyclin genes, we turned our attention to TRF2-interacting proteins. TRF2 was recently shown to interact with M1BP (motif 1 binding protein) [43]. Interestingly, M1BP was recently demonstrated to interact with GFZF, a nuclear glutathione S-transferase protein that has been implicated in cell cycle regulation [44]. We suspected that GFZF may interact with TRF2. Indeed, using FLAG immuno-affinity purification from FLAG-HA TRF2-inducible S2R+ cells followed by mass spectrometry analysis, we discovered that both M1BP and GFZF are in complex with the evolutionarily conserved TRF2 (also known as short TRF2), but not with the long *Drosophila*-only TRF2 isoform or with TBP (Table 3 and Table S3). This prompted us to examine the occupancy of TRF2, M1BP and GFZF in the vicinity of the TSSs (−100 to +100 relative to the TSS) of TRF2-regulated cyclin genes, using publicly available TRF2, M1BP and GFZF ChIP-exo analyses in *Drosophila* S2R+ cells (GSE97841, GSE105009) [43, 44]. We examined the number of bound sites, the average peak scores and the maximum peak scores of cyclin genes and several ribosomal protein genes for comparison (Table 4). As expected, M1BP, TRF2 and GFZF co-occupy the promoters of the ribosomal protein genes. Interestingly, while M1BP occupies the −100 to +100 regions of all the examined cyclin genes, both TRF2 and GFZF occupy the −100 to +100 regions of *Cyc A, Cyc B, Cyc B3* and *Cyc E*, and to a lesser extent the promoters of *Cyc C* and *Cyc D*, which are not regulated by TRF2. The occupancies of the three proteins is especially striking in the vicinities of the *Cyc B* and *Cyc B3* TSSs. Thus, the effects of TRF2 knockdown on cyclin gene expression generally correlate with the occupancies of both TRF2 and GFZF in the −100 to +100 regions of *Cyc A, Cyc B, Cyc B3* and *Cyc E*.

To characterize the co-occupancies of TRF2, GFZF and M1BP in a genomewide manner, we examined the binding of each factor to *Drosophila* promoter regions (± 50 bp relative to FlyBase annotated TSSs). Remarkably, a major fraction of promoters is bound by all three transcription factors (Fig. 5a). Reassuringly, the co-bound promoters include the TRF2-regulated *Cyc A, Cyc B, Cyc B3* and *Cyc E*, but not *Cyc C* and *Cyc D* promoters, which are not regulated by TRF2.

**Fig. 5.**
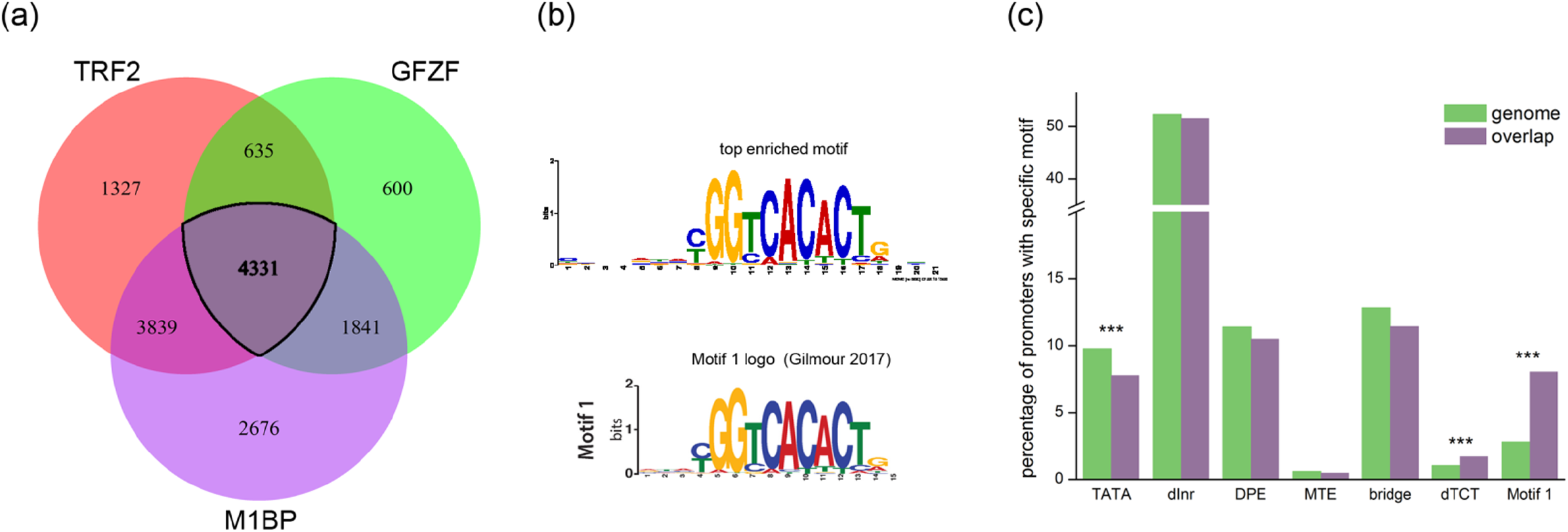
Promoters of genes bound by TRF2, GFZF and M1BP are enriched for specific core promoter motifs. (a) Schematic representation of genes containing at least one binding site of the specified transcription factor at ±50bp relative to its TSS, as determined by FlyBase. ChIP-exo data was retrieved from GSE97841 and GSE105009. (b) Top enriched motif among the 4331 commonly bound promoters, as detected by MEME analysis. Its resemblance to Ohler Motif 1 is depicted by the motif logo derived by M1BP ChIP-exo [43]. (c) The 4331 commonly bound regions were analyzed for core promoter composition. This promoter group was found to be depleted for the TATA-box motif and enriched for dTCT and the Motif 1 core promoter elements. p-values were adjusted using Bonferroni correction. ***p< 10^-5^.

To better decipher the characteristics of the co-bound promoters, we used MEME [45] to detect enriched sequence motifs. Interestingly, the top enriched motif in promoters that are bound by all three proteins (Fig. 5b) closely resembles Ohler Motif 1 [46], also detected in M1BP ChIP-exo analysis [43]. We next analyzed the core promoter composition of the co-bound promoters, using the ElemeNT algorithm [47]. Strikingly, the co-bound promoters are depleted for the TATA-box motif and enriched for the TCT and Motif 1 core promoter elements, as compared to the genomic distribution of core promoter elements (Fig. 5c).

To examine the contribution of M1BP and GFZF to the effect of TRF2 knockdown on cyclin gene expression, we used RNAi to knockdown the expression of either M1BP, GFZF, TRF2 or their combinations. The use of each of the dsRNA probes resulted in a significantly reduced expression of the targeted gene (Fig. 6a). Surprisingly, M1BP knockdown resulted in increased expression of *Trf2, gfzf* and *Cyc E*. Since both TRF2 and M1BP were previously shown to affect the expression of ribosomal protein genes [32, 48], we tested whether their knockdown affects the expression of several ribosomal target genes, namely, *RpL30, RpLP1* and *RpLP2*. While TRF2 and GFZF knockdown did not affect their expression, M1BP knockdown resulted in significantly increased expression of *RpLP2* (Fig. S5a). Notably, this effect was not general, but rather specific to distinct cyclin and ribosomal protein genes (Fig. 6 and Fig. S5), as the expression of *CG12493* and *Sgll*, two previously identified M1BP targets [48], was reduced following M1BP knockdown (Fig. S5b). The expression levels of *Cyc A* and *Cyc B* were specifically reduced following TRF2 knockdown, but not following GFZF or M1BP knockdown (Fig. 3a and Fig. 6). *Cyc D* expression was not affected by either of these single factor knockdowns (Fig. 6a), as in Fig. 3. As can be observed by TRF2 knockdown, as well as by the combined knockdown of TRF2, GFZF and M1BP, the expression pattern of *Cyc A* and *Cyc B* are mostly influenced by TRF2 knockdown. *Cyc E* exhibits a composite pattern: it is reduced following TRF2 knockdown, but the combined knockdowns of TRF2 and M1BP or GFZF seem to restore its expression, as compared to mock treatment. Notably, *Cyc D* expression pattern is the least affected by the different knockdown combinations.

**Fig. 6.**
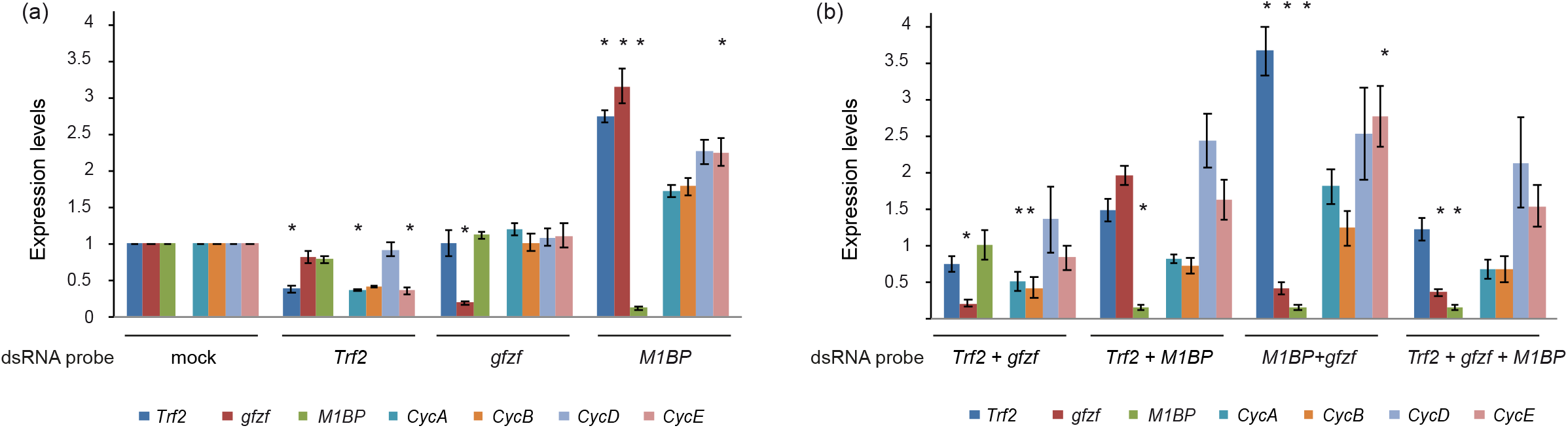
*Cyclins A* and *B* expression levels are mostly affected by TRF2 knockdown, and to a lesser extent by GFZF and M1BP. *Drosophila* S2R+ cells were incubated for three days with dsRNA probes directed against *Trf2* (probe #1), *gfzf, M1BP* or their combinations. RNA was isolated from the cells and reverse transcribed (RT) to cDNA. Real-time PCR (qPCR) experiments were used to analyze the RNA levels of the endogenous of *Trf2, gfzf* and *M1BP*, as well as *CycA, CycB, CycD* and *CycE* genes, as indicated. (a) Single knockdowns of *Trf2, gfzf* or *M1BP*. (b) Knockdown of multiple genes, as indicated. qPCR experiments were performed in triplicates, and the graph represents the average of 4 independent experiments. Error bars represent the SEM. *p< 0.05, one-way ANOVA followed by Tukey’s post hoc test as compared to the mock treatment of the relevant gene.

Taken together, these data suggest that the observed effects of TRF2 knockdown on cell cycle progression (Figs. 1, 2 and 4) are, at least partially, mediated by the reduced expression of *Cyc E, Cyc A* and *Cyc B* following TRF2 knockdown (Figs. 3 and 6). Importantly, the co-occupancy of TRF2, GFZF and M1BP in the promoters of these cyclin genes, the enrichment of Motif 1 in their promoters and the expression patterns of the cyclin genes following the knockdown of either TRF2 alone, or in combination with GFZF and/or M1BP, imply that GFZF and M1BP may serve as co-factors in TRF2-regulated cell cycle progression. Yet, it is TRF2 that is the major transcription factor regulating the expression pattern of the abovementioned cyclin genes.

## Discussion

In this study, we discovered that knockdown of the general/basal transcription factor TRF2 results in accumulation of cells in G1 and in reduction in the number of cells in S and G2/M phases. G1/S transition is regulated by Cyclin E activity, while S phase and G2/M transition are regulated by Cyclin A activity, and transition into and within mitosis is regulated by the activities of Cyclin B and Cyclin B3. Remarkably, TRF2 knockdown in S2R+ cells resulted in reduced expression of *Cyc E, Cyc A, Cyc B* and *Cyc B3* (but not *Cyc D)*, suggesting that TRF2 regulates cell cycle progression by modifying the expression of specific cyclins. The reduced expression of *Cyc A, Cyc B* and *Cyc B3* and the reduction of the number of cells undergoing mitosis following TRF2 knockdown, are in line with previous studies, which demonstrated inhibition of nuclear mitotic entry in *Drosophila* embryos following simultaneous knockdown of *Cyc A, Cyc B* and *Cyc B3* [49]. Interestingly, Cyclin A and Cyclin B have previously been shown to inhibit metaphase-anaphase transition, whereas Cyclin B3 promotes it [50]. Thus, the specific accumulation of cells in anaphase and telophase observed by imaging flow cytometry (Fig. 4e, f), could result from the reduced expression of *Cyc A* and *Cyc B* following TRF2 knockdown (Fig. 3). Notably, to the best of our knowledge, this study is the first to employ imaging-flow cytometry in the analysis of *Drosophila* cells.

To examine whether TRF2-interacting proteins could account for the observed regulation of *Cyc E, Cyc A, Cyc B* and *Cyc B3* (but not *Cyc D* or *Cyc C*), we searched for TRF2-interacting proteins. TRF2 has been shown to interact with M1BP [43], and M1BP has been shown to interact with GFZF, a nuclear glutathione S-transferase implicated in cell cycle regulation [43]. Our proteomic analyses revealed that both M1BP and GFZF preferentially interact with TRF2, but not with TBP (Table 3 and Table S3). Remarkably, examination of publicly available TRF2, M1BP and GFZF ChIP-exo data from *Drosophila* S2R+ cells [43, 44], indicated that the effects of TRF2 knockdown on the expression of cyclin genes correlate with TRF2, GFZF and M1BP co-occupancies of the promoters of the TRF2-regulated cyclin genes (Table 4), in line with the reported regulatory effects of GFZF. Genome-wide examination of promoters bound by all three transcription factors revealed the enrichment of the TCT and Motif 1 core promoter elements. Whereas one would expect the TCT to be enriched as TRF2 and M1BP have previously been implicated in the regulation of ribosomal protein genes [32], the enrichment of Motif 1 within promoters bound by all three factors indicates a shared function for TRF2, GFZF and M1BP. Notably, in our experimental system, TRF2 knockdown by RNAi resulted in a two-fold reduction in *Trf2* levels (Fig. 3a, Table S1). Under these conditions, we did not detect any change in ribosomal protein gene expression (Table S1). Hence, the reduced expression of *Cyc A, Cyc B* and *Cyc E* following a two-fold reduction in TRF2 expression, does not result from a general inhibition of protein synthesis. As direct binding of TRF2 to DNA could not be demonstrated [32], it is likely that TRF2 indirectly regulates the expression of these cyclin genes and that there are TRF2-associated factors that enable DNA binding. Our analysis suggested that GFZF and M1BP could serve as such factors for specific TRF2-regulated processes. Interestingly, the expression patterns of the cyclin genes following the knockdown of either TRF2 alone, or in combination with GFZF and/or M1BP, indicated that among these three factors, TRF2 is the major contributor to the expression pattern of *Cyc A, Cyc B* and *Cyc E*. Moreover, while *Cyclin A, Cyclin B* and *Cyclin E* expression levels are reduced following TRF2 (but not GFZF) knockdown, the expression levels of both *Cyclin A* and *Cyclin B*, but not *Cyclin E*, are reduced following the combined knockdown of TRF2 and GFZF (Fig. 6), suggesting that the reduced expression levels of *Cyclin A* and *Cyclin B* are not mediated via *Cyclin E*.

Notably, mouse TBP was recently shown to remain bound to mitotic chromosomes during mitosis of mouse embryonic stem cells (mESCs), and to recruit a small population of Pol II molecules to mitotic chromosomes [51]. Nevertheless, active Pol II transcription occurs in the absence of mouse TBP, whereas Pol I and Pol III, are significantly reduced [11, 51]. It remains to be determined whether *Drosophila* TBP is bound to mitotic chromosomes during mitosis. As we did not observe significant effects on cell cycle progression or mitosis following *Drosophila* TBP knockdown (Figs. 1, 2 and 4), it is likely that *mouse* TBP may exert different functions as compared to *Drosophila* TBP, perhaps via its associated proteins.

The effects of TRF2 knockdown on cell cycle progression of *Drosophila* cultured S2R+ cells are in line with the defects observed in embryonic development of TRF2 knockout flies. TRF2 has also been shown to be essential for embryonic development of *C. elegans*, zebrafish and *Xenopus*. It remains to be determined whether knockdown of TRF2 in cellular systems from these species results in similar effects.

Taken together, using *Drosophila* cells as a model system, we discovered that the knockdown of TRF2, rather than TBP, regulates cell cycle progression via distinct target genes. Importantly, we discovered that TRF2 is associated with the GFZF and M1BP proteins, and that TRF2, GFZF and M1BP co-occupy the promoters of the TRF2-regulated cyclins. Furthermore, we show that TRF2, rather than GFZF or M1 BP, is the major contributor to the expression pattern of *Cyc E, Cyc A* and *Cyc B*. Importantly, while a general transcription factor may be regarded as having a “generic function”, our findings emphasize the unique, unanticipated functions of *Drosophila* TRF2 as an essential factor for specific major cellular processes.

## Materials and Methods

### *Drosophila melanogaster* Schneider S2R+ Cells

*Drosophila melanogaster* Schneider S2R+ adherent cells were cultured in Schneider’s *Drosophila* Media (Biological Industries) that was supplemented with 10% heat-inactivated FBS and Penicillin 100 units/ml Streptomycin 0.1mg/ml (Biological Industries).

### Generation of dsRNA probes

All dsRNA probes were chosen based on http://www.dkfz.de/signaling/e-rnai3// and http://www.flyrnai.org/snapdragon as described in [52]. Primer sequences used for the generation of dsRNA probes are provided in Table 1. DNA fragments corresponding to each dsRNA were subcloned into both pBlueScript SK+ and KS+. The dsRNA probes were generated by PCR amplification of the DNA using T7 and T3 primers, followed by *in vitro* transcription of templates in both pBlueScript orientations using T7 RNA polymerase. Resulting RNA products were annealed to generate the dsRNA probes.

**Table 1.**
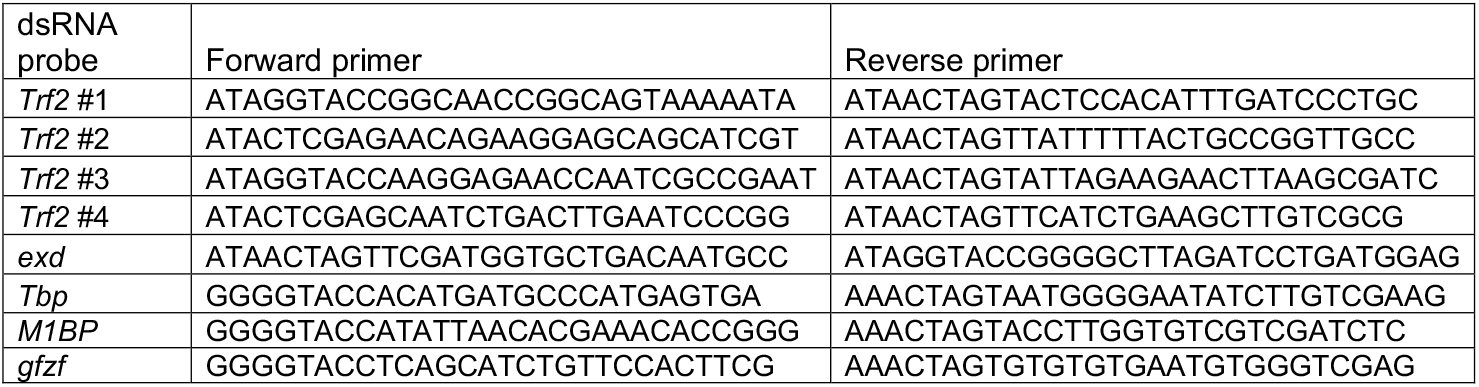
Primers for generation of dsRNA probes

### RNA interference (RNAi)

For 6 well plate, 1.25×10^6^ cells/well were resuspended and seeded in empty Schneider’s *Drosophila* Media (Biological Industries) with 30μg/ml dsRNA directed against different genes for 1 hour. Next, two volumes of complete medium were added to the wells, and cells were incubated for 3 additional days.

### Western blot analysis

Knockdown of TRF2 and TBP was verified by western blot analysis using anti-TRF2 and anti-TBP polyclonal antibodies (generous gift from Jim Kadonaga). Cyclin E levels were analyzed by the 8B10 antibodies (generous gift from Helena Richardson) [53]. The levels of Actin or γ-Tubulin, as a loading control, were detected using either mouse monoclonal anti-Actin (Abcam, 8224) or anti-γ-Tubulin (Sigma, GTU-88) antibodies. Anti-Cyclin A and -Cyclin B concentrated monoclonal antibodies (Developmental Studies Hybridoma Bank, A12 and F2F4, respectively) were tested as well, however no endogenous proteins were detected, possibly due to technical limitations.

### RNAi-coupled overexpression

For 6 well plate, 1.25×10^6^ *Drosophila* S2R+ cells/well were resuspended and seeded in empty medium with 30μg/ml dsRNA directed against TRF2 for 1 hour. Next, two volumes of complete medium were added to the wells and cells were incubated for 3 days. Three days post dsRNA treatment, cells were transfected with the TBP-pAc expression vector (930 ng) or an empty pAc vector control using the Escort IV reagent (Sigma). Media was replaced 18-24 hrs post transfection. Cells were harvested 36-48 hrs post transfection, and RNA was purified and analyzed by RT-qPCR. Each qPCR experiment was performed in triplicates. The graphs represent an average of 3 independent experiments. Error bars represent SEM.

### RNA-seq analysis

S2R+ cells were treated with dsRNA probes against *Trf2* (probe #1) and *Tbp*, and harvested at two time points - 48h and 72h post RNAi treatment. For each time point, a matched mock control was collected separately.

RNA was extracted using Quick-RNA™ MiniPrep (Zymo Research), and 800ng of each sample was purified using NEBNext Poly(A) mRNA Magnetic Isolation Module (NEB #E7490). Libraries were prepared using NEBNext Ultra II RNA Library Prep Kit for Illumina (NEB #E7770), following the manufacturer’s instructions. NEBNext Multiplex Oligos for Illumina (NEB #E7335, NEB #E7500, NEB #E7710, NEB #E7730) were used. Libraries were pooled and a 1% PhiX library control was added. Single-end sequencing was performed on an Illumina NextSeq 500 machine.

Reads were aligned to dm6 genome build using STAR (version 2.6.0a), and htseq-count (version 0.5.1p3, [54]) was used to count the reads mapped to each gene. Differential expression analysis of conditions was performed using the DESeq2 R package [55]. Only genes with adjusted p-value < 0.1 were considered for subsequent analysis. GO terms analysis was carried out using STRING v11 [56]. For all experiments, three independent biological replicates were compared and merged for subsequent analysis. RNA-seq Data is available at GSE133685.

### RT-PCR

Total RNA was isolated using the PerfectPure RNA Cultured Cell kit (5 PRIME) or Quick-RNA™ MiniPrep (Zymo Research). One microgram of the total RNA was reverse-transcribed into cDNA with M-MLV (Promega) or qScript Flex cDNA Kit (Quanta). Control reactions lacking reverse transcriptase were also performed to ensure that the levels of contaminating genomic DNA were negligible. Quantitation was performed by real-time PCR to determine the transcription levels of the endogenous genes. The expression levels were compared to *Gapdh2*. Primer sequences for real-time PCR are provided in Table 2. For all quantifications, the error bars represent ±S.E.M of at least 3 independent experiments; NS, not significant; \*\**P*<0.01; \*\*\**P*< 0.001. Statistical analyses were performed on log-transformed relative quantification (RQ) values using one-way ANOVA followed by Tukey’s post hoc test, unless otherwise stated in the figure legend.

**Table 2.**
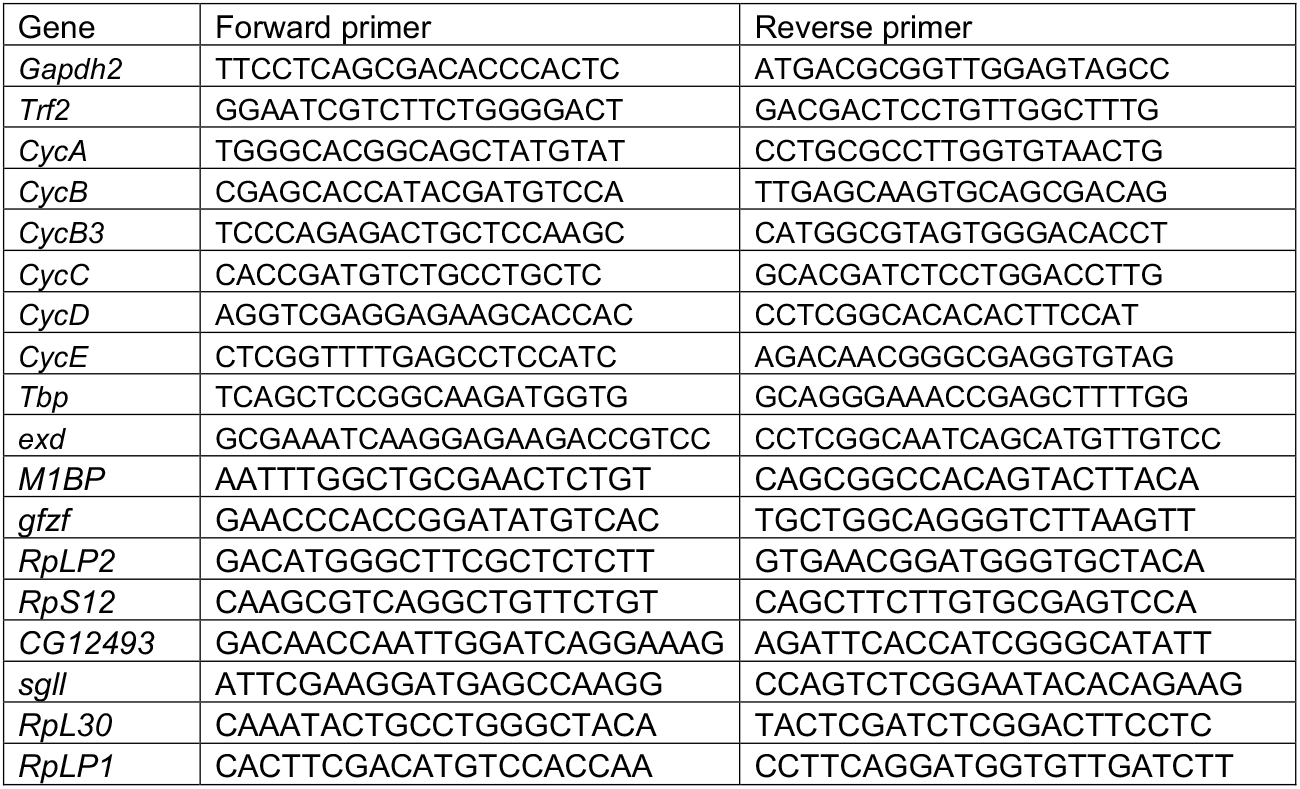
Real-time PCR (qPCR) primers

**Table 3.** Enrichment of TRF2, TBP, M1BP and GFZF in FLAG-immuno-affinity purified complexes from inducible S2R+ cells

| Name of purified (/co-purified) protein | Enrichment of purified (/co-purified) proteins within |  |  |
| --- | --- | --- | --- |
|  | Short TRF2-associated proteins | Long TRF2-associated proteins | TBP-associated proteins |
| TRF2 | 78.695 | 6.128 | - |
| TBP | - | - | 18.3 |
| M1BP | 1.795E7 | - | - |
| GFZF | 1.683E7 | - | - |

The values in the table represent the enrichment of each purified (/co-purified) protein in the indicated sample. The enrichment was calculated as the ratio between the mass spectrometry area (the average of the three most intense peptides from each protein) of the induced sample and the un-induced sample.

**Table 4.**
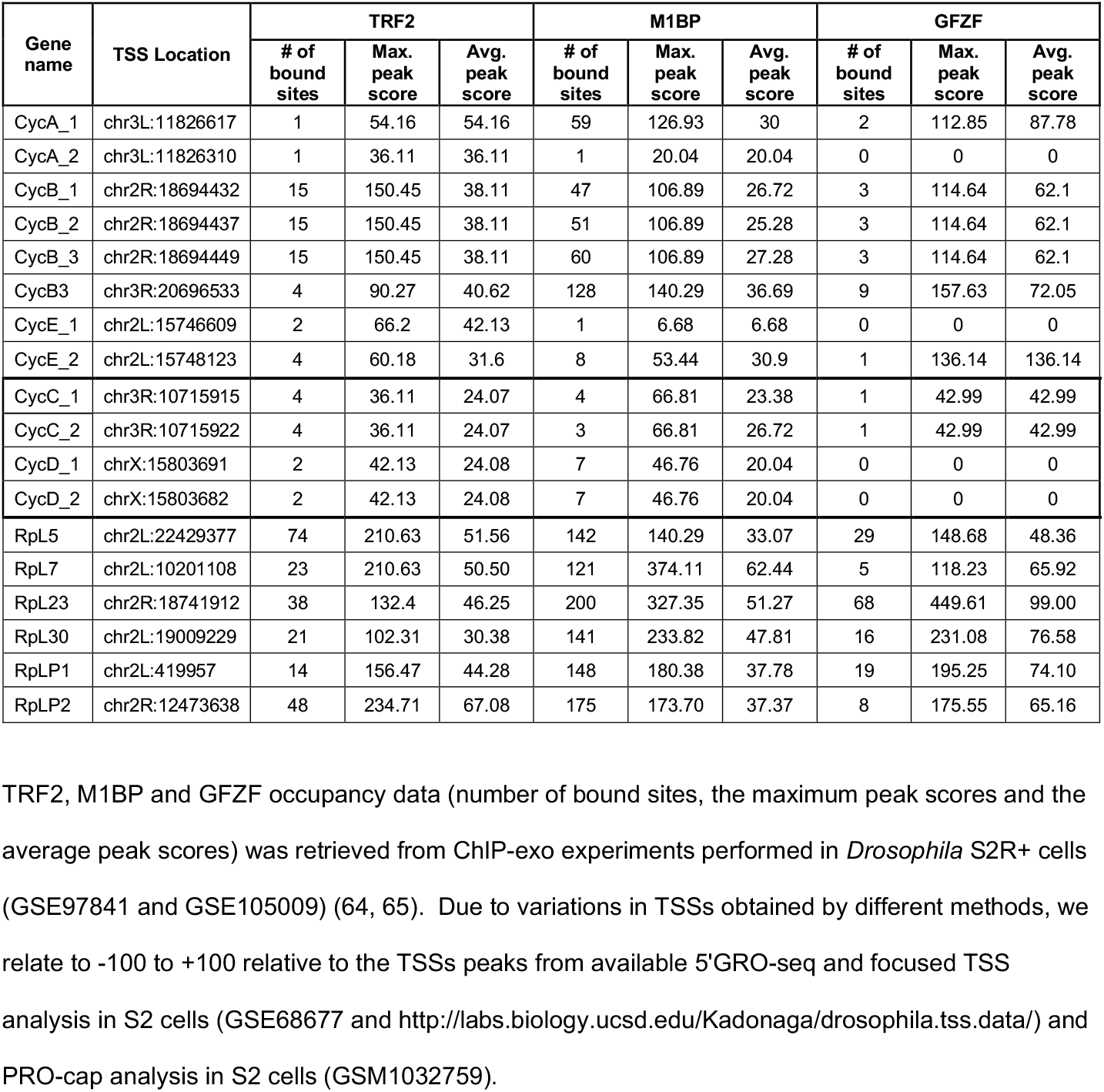
TRF2, M1BP and GFZF occupancy within −100 to +100 relative to the TSSs

### Flow cytometry analysis

For cell cycle distribution by Propidium Iodide (PI) staining, cells were harvested following 72h incubation with dsRNA, centrifuged for 5 min at 300g and fixed with 80% ethanol at 4°C overnight. Before subjecting the cells to flow cytometry, the cells were centrifuged for 5 min at 300g, washed in 1 ml of Phosphate-buffered saline (PBS) and incubated for 40 min at 4°C. The cells were then stained in PBS containing 50 μg/ml PI (Sigma) and 50 μg/ml RNase A (Roche). After incubation for 15 min at room temperature, fluorescence was measured using a FACSCalibur Becton Dickinson flow cytometer.

For G1 phase cell arrest by hydroxyurea (HU) and BrdU (5-Bromo-2’-Deoxyuridine)-PI staining, 72h following incubation with dsRNA, the medium was replaced with medium containing a final concentration of 1mM HU for 18h [57] (for control cells, the medium was replaced with a fresh medium). BrdU (40μM final concentration) was added to the medium for 2h. Cells were released from HU by medium replacement and, at 0, 2, 4, 6 or 8 hours following the release, cells were harvested, centrifuged and fixed with 80% ethanol at 4°C overnight. Following fixation, cells were stained with FITC (*Fluorescein isothiocyanate*)-conjugated anti-Brdu antibodies (BD) and PI according to the provided protocol, and fluorescence was measured using BD FACSARIA III. All flow cytometry data was analyzed using the FlowJo software. Statistical analyses of flow cytometry and imaging flow cytometry data were performed in SPSS using two-tailed Students t-test. The number of times each experiment was repeated, is detailed in the figure legends.

It is of note that unfortunately, we were unable to synchronize cells in G2/M in a reversible manner using either Nocodazole or Colchicine, which cause microtubules depolymerization. Specifically, Nocodazole did not arrest the S2R+ cells in G2/M, while Colchicine, which has been used since the 1950s to inhibit mitotic progression, did cause enrichment of mitotic cells (Fig. 4b, d). However, this effect was irreversible (data not shown). Thus, Colchicine could not be used for G2/M block-release synchronization experiments.

### Immunostaining for fluorescence microscopy and Imaging flow cytometry analysis

Cells were RNAi-treated as described above. On day 4, 2ml of fresh medium was added to the wells. To enrich for G2/M, as a control, Colchicine (Sigma) was added to a final concentration of 350ng/ml. On the following day, the cells were harvested and fixed with 4% Paraformaldehyde (PFA)/PBS (30 min), washed in PBST (PBS containing 0.5% Triton X-100), blocked with PBS containing 1% BSA and 1% serum (1h), and incubated with phospho-Histone H3 (Ser10) antibody (1:200, Cell Signaling Technology #9701) for 1h at RT, followed by overnight at 4°C. Cells were washed, stained with the secondary antibody (1:500, DyLight 488, ab96883), and then counter-stained with 10μg/ml Hoechst 33342 (Sigma). For microscopy analysis, samples were also stained for filamentous Actin with 3.5μM Acti-stain 670 Phalloidin (Cytoskeleton, Inc. Cat. # PHDN1). Following staining, samples were subjected to imaging flow cytometry, confocal microscopy or wide-field fluorescence microscopy analysis.

### Microscope image analyses

High resolution images were acquired using a Leica SP8 confocal microscope, and high-throughput images for quantitative analysis were acquired using a Leica DMi8 microscope. Three separate experiments were performed and captured at 20x magnification. For each treatment, approximately 275 frames were acquired and analysed. The total number of Alexa 488 anti-phosphor-Histone H3 (Ser10) (PH3) labeled cells, and the total number of Hoechst stained cells, were calculated using the Fiji distribution of ImageJ.

Analysis workflow:

1. The raw PH3 channel images were enhanced using brightness and contrast, and then the background was subtracted by reducing the Gaussian blurred filtered image of the enhanced image. Next, a median filter was applied to smoothen the image and an Otsu threshold was applied to get the binary image of all mitotic nuclei. Finally, watershed was implemented to separate touching nuclei. Mitotic cells were counted, eliminating small debris and noise.
2. To analyze the Hoechst channel, the background was subtracted by reducing the Gaussian blurred filtered image of the original image. Next, a Moments threshold was applied to get the binary image of total nuclei. Finally, watershed was implemented to separate touching nuclei. Nuclei were counted while eliminating small debris and noise.

The ratio between the number of mitotic cells and the total number of cells yields the mitotic index for each treatment.

All manipulations in the images were made evenly across the entire field. The Fiji macros will be shared upon request.

### Multispectral imaging flow-cytometry (IFC) analysis

Cells were imaged using multispectral imaging flow cytometry (ImageStreamX mark II imaging flow-cytometer; Amnis Corp, Seattle, WA, Part of EMD Millipore). Each experiment was performed 3 times. In each experiment, at least 40,000 cells were collected from each sample, and data were analyzed using the image analysis software (IDEAS 6.2; Amnis Corp). Images were compensated for fluorescent dye overlap by using single-stain controls. Imaging flow cytometry results were analyzed by calculation of a set of parameters, termed “features”, performed on a defined area of interest, termed “mask”. The serial gating strategy to identify the mitotic cell population was as follows: Single cells were first gated using the area and aspect ratio features on the bright-field (BF) image (the aspect ratio, which indicates how round or oblong an object is, is calculated by division of the minor axis by the major axis). Uncropped cells were gated using the centroid X (the number of pixels in the horizontal axis from the upper left corner of the image to the center of the mask) and area features. Focused cells were gated using the Gradient RMS feature, as previously described [58] (Fig. S3a-c). Following this standard gating series, the mitotic cell fraction of the entire cell population was identified using the staining intensity for PH3 AF488 (channel 2), and mitotic cells were gated as the high intensity population of PH3 staining within all focused cells (Fig. S3d, e).

For a more complex analysis, we performed a second gating series. Focused cells were first gated for G2/M based on DNA (Hoechst) intensity (Fig. S3) and then gated for mitotic cells, as previously, by high PH3 intensity. To further subdivide into the specific cell division phases, we gated according to nuclear morphology based on the spot count and aspect ratio intensity features (see Fig. S3f, g for detailed masking and gating). As it was previously shown that serine 10 of histone H3 becomes dephosphorylated during telophase [59], the telophase population was derived from the negative PH3-stained cells, based on the BF circularity feature and DNA aspect ratio intensity (see Fig. S3h for detailed masking and gating). Full details of all masking, features and analysis strategies are included in the legend of Fig. S3.

### Identification of unique TRF2-interacting proteins

To identify the proteins that are in complex with TRF2 (the evolutionarily conserved short TRF2), we used inducible FLAG-HA-tagged TRF2 S2R+ cells [52]. As controls, we used inducible S2R+ for FLAG-HA-long TRF2 [52] or FLAG-HA-TBP (generated as in [52]). Cells were either induced by copper sulfate or left untreated. Protein extracts were prepared and TRF2-containing complexes were immunoprecipitated using anti-FLAG M2 affinity gel (Sigma). Following the IP, TRF2-containing complexes were released with a FLAG peptide (Sigma). Samples were resolved by SDS-PAGE. Proteins that were purified from TRF2-induced cells were separated into two samples: proteins larger or smaller than 40 kDa. Samples were subjected to Mass spectrometry analyses (The Smoler Protein Research Center, Technion). Briefly, samples were digested by trypsin, analyzed by LC-MS/MS on Q-Exactive Plus (ThermoFisher) and identified by the Discoverer software (with two search algorithms: Sequest (ThermoFisher) and Mascot (Matrix science) against the

*Drosophila melanogaster* section of the NCBI non-redundant and Uniprot databases, and a decoy database (in order to determine the false discovery rate). All the identified peptides were filtered with high confidence, top rank, mass accuracy, and a minimum of 2 peptides. High confidence peptides have passed the 1% FDR threshold. Semi-quantitation was done by calculating the peak area of each peptide. The area of the protein is the average of the three most intense peptides from each protein. The results are provided in Table S3.

### Visualization of publicly available ChIP-exo data

A genome browser session (https://genome.ucsc.edu/s/Anna%20Sloutskin/dm3_ChIP_Exo) based on available TRF2, GFZF and M1BP ChIP-exo bedgraph files (GSE97841, GSE105009) [43, 44] was created. The session contains an “Overlap” track (the ChIP-exo peaks that were identified as overlapping in the ±50bp window relative to FlyBase TSS) and the “trustedTSS” track that is based on 5’ GRO-seq (GSE68677) and PRO-Cap (GSM1032759) data. The relevant interval (±50bp or 100bp) is indicated.

The authors declare that they have no competing interests

## Abbreviations

ChIP-exo: chromatin immunoprecipitation combined with exonuclease digestion followed by high-throughput sequencing
DPE: downstream core promoter element
GFZF: GST-containing FLYWCH zinc-finger protein
M1BP: Motif 1 binding protein
Pol II: RNA polymerase II
TBP: TATA-box-binding protein
TRF2: TBP-related factor 2
TSS: Transcription start site

## Authors’ contributions

A. Kedmi prepared RNA samples and O. Yaron and A. Sloutskin performed RNA-seq experiments. T. Doniger and A. Sloutskin analyzed the RNA-seq experiments and performed bioinformatics analysis. A. Kedmi, N. Epstein, L. Gasri-Plotnitsky and D. Ickowicz performed and/or analyzed flow cytometry experiments. A. Kedmi, A. Sloutskin and D. Ideses performed reverse transcription-qPCR analysis. I. Shoval and Z. Porat advised and performed the analysis of ImageStream^®^ experiments, A. Kedmi and I. Shoval performed and analyzed fluorescence microscopy experiments. E. Darmon and D. Ideses performed western blot analyses. A. Sloutskin performed the statistical analysis. A. Kedmi, A. Sloutskin and T. Juven-Gershon designed the study, planned experiments, analyzed results, and wrote the manuscript with input from all authors.

## Acknowledgements

We thank Jim Kadonaga for the generous gift of anti-TRF2 and anti-TBP antibodies, for invaluable advice and assistance and for critical reading of the manuscript. We thank Helena Richardson for the generous gift of anti-Cyclin E antibodies. We thank Moshe Oren, Doron Ginsberg, Ze’ev Paroush, Sivan Henis-Korenblit, Shaked Cohen and Itay Lazar for invaluable advice and assistance. We thank Dr. Jennifer Israel-Cohen for assistance with statistical analysis. We thank Doron Ginsberg, Yaron Shav-Tal, Galit Shohat-Ophir and Orit Adato for critical reading of the manuscript. We thank Orit Adato and Sarit Lampert for technical assistance.

Supplementary data is provided in a separate PDF (Figures S1-S5, Table S2) or Excel Files (Table S1, Table S3).

