## Supplementary material for "The TRF2 General Transcription Factor Is a Key Regulator of Cell Cycle Progression": Figures S1-S5, Table S2

### Is a Key Regulator of Cell Cycle Progression

#### SUPPLEMENTAL INFORMATION

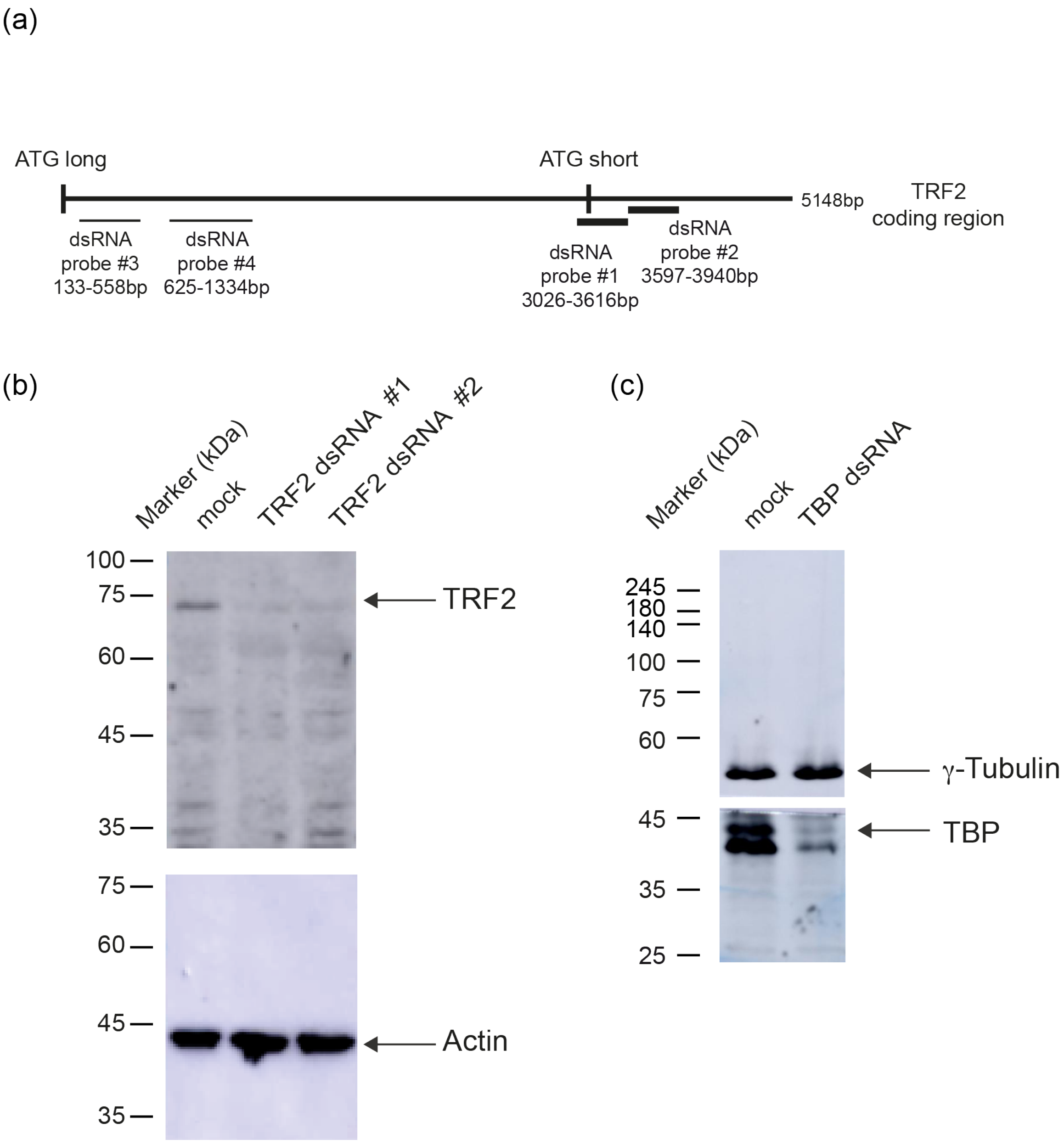


**Fig. S1.** (a) Schematic representation of dsRNA TRF2 probes used in this study. (b) Western blot analysis following TRF2 and (c) TBP knockdown in S2R+ cells, using anti-TRF2 and anti-TBP polyclonal antibodies, respectively. Actin and γ-Tubulin were used as loading controls.

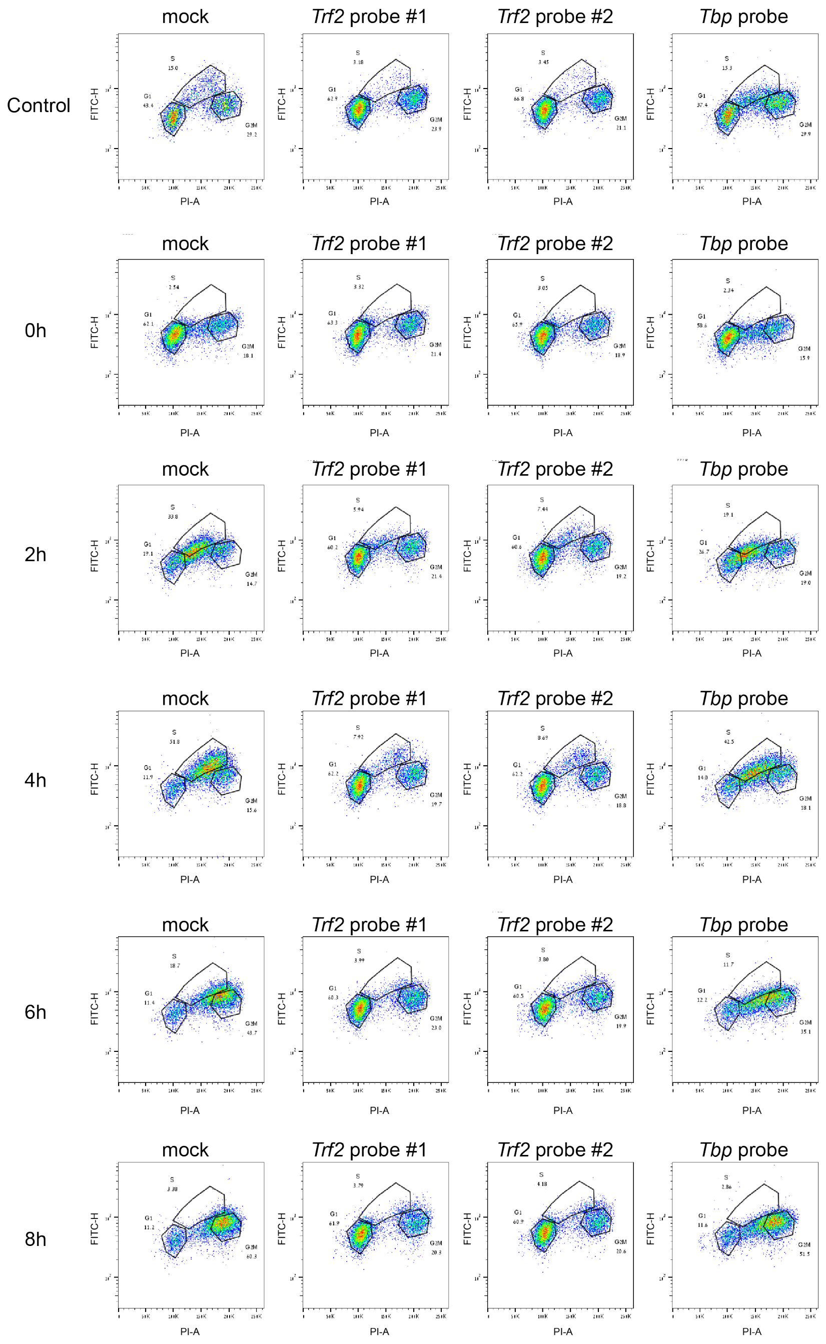


**Fig. S2.** TRF2 is involved in S phase progression. *Drosophila* S2R+ cells were incubated for three days with dsRNA probes directed against *Trf2* or *Tbp*. Next, cells were either left untreated or treated with 1mM Hydroxyurea for 18h. The cells were allowed to resume cell cycle for 2h, 4h, 6h or 8h in fresh medium containing 40µM BrdU, and were then fixed with 80% ethanol overnight and analyzed by FACS using BrdU-PI staining. The PI fluorescence intensity (representing DNA content) is plotted on the X-axis (linear scale), and the BrdU-FITC fluorescence intensity (representing BrdU incorporation into the DNA) is plotted on the Y-axis (log scale).

Kedmi et al._Fig. S3

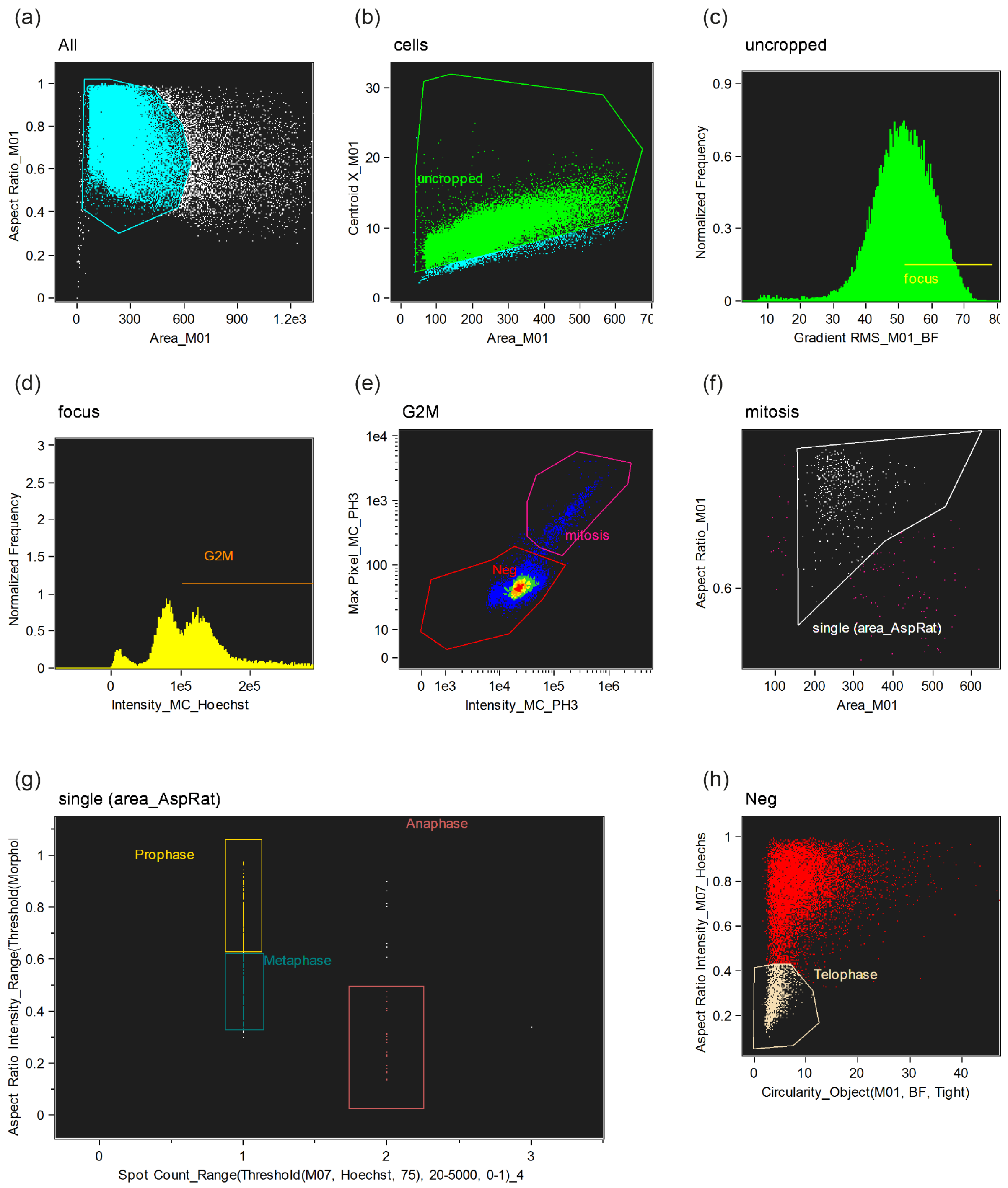


**Fig. S3.** Gating and masking strategy for imaging-flow cytometry data analysis. (a) Cells were gated for single cells, using the area and aspect ratio features on the BF image. (b) Centered and uncropped cells were gated based on the centroid X (the number of pixels in the horizontal axis from the upper left corner of the image to the center of the mask) and area features. (c) Focused cells were gated, using the Gradient RMS feature, as previously described [1]. (d) The G2/M population was gated out of the focused cells, based on DNA (Hoechst) intensity. (e) Mitotic cells were gated from the G2M population, as the high intensity population of pH3 staining, based on

the intensity feature of pH3 and Max pixel feature of pH3 (the largest value of the background-subtracted pixels contained in the input mask). (f) To include only single positive pH3 stained cells, doublet cells were eliminated by gating the single cells from the mitotic population, using the area and aspect ratio features of the BF. (g) To discriminate between the different mitotic phases subpopulations, several masks were created: 1. A morphology mask that includes all pixels within the outermost image contour. 2. A threshold mask that includes the highest intensity pixels (indicated as percentages). 3. A range mask that selects components in an image within a selected size (μm), was used to eliminate small components.
Furthermore, the following features were used on the combined masks:
1. Spot count - the number of connected components in an image.
2. Aspect ratio intensity - the aspect ratio weighted for fluorescence intensity.
These masks were combined and the features were calculated and plotted as follows: Spot count of the combined mask: range (threshold 75%, M07) 20-5000, was plotted against the aspect ratio intensity of the combined mask: range (Threshold(Morphology(M07), 82%) 15-5000.
The prophase population was defined as having a more circular nuclear staining (aspect ratio intensity should be high), and was hence gated as one nuclear spot count with aspect ratio intensity bigger than 0.6. On the other hand, the metaphase population was defined as having a more elongated DNA distribution and was gated as one nuclear spot count with aspect ratio intensity smaller than 0.6. Finally, the anaphase population was gated as cells with two nuclear spots having aspect ratio intensity less than 0.6 [2]. (h) As serine 10 of histone H3 becomes dephosphorylated during telophase, the telophase population was derived from the negative pH3-stained cells. To identify telophase pairs, an object mask was created (which segments images to closely identify the area corresponding to the cell). The circularity feature (which measures the degree of the mask's deviation from a circle) of the object mask (BF), was plotted against the aspect ratio intensity of the M07 DNA mask. Telophase cells were gated as having the lowest BF circularity and as the most elongated, based on DNA stain (lowest aspect ratio intensity (M07)).

(a)

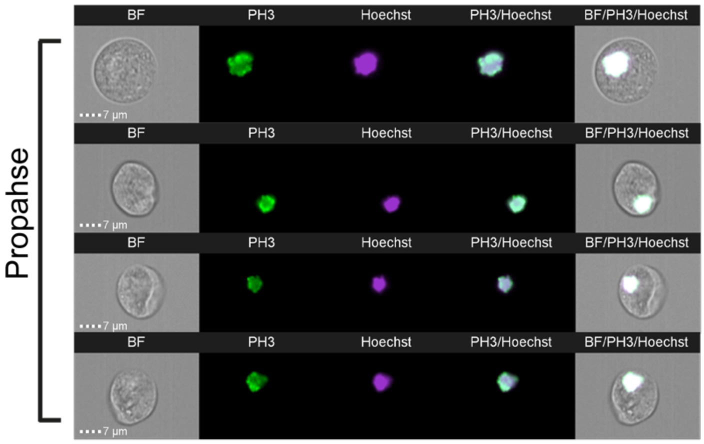


(b)

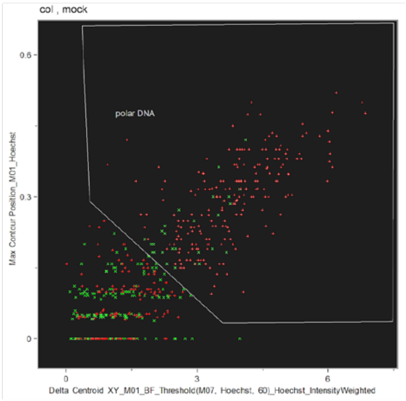


| Population | Count | %Gated |
|---|---|---|
| Colchicine treatment | 339 | 100 |
| Polar DNA | 210 | 61.9 |
| mock | 190 | 100 |
| Polar DNA | 27 | 14.2 |

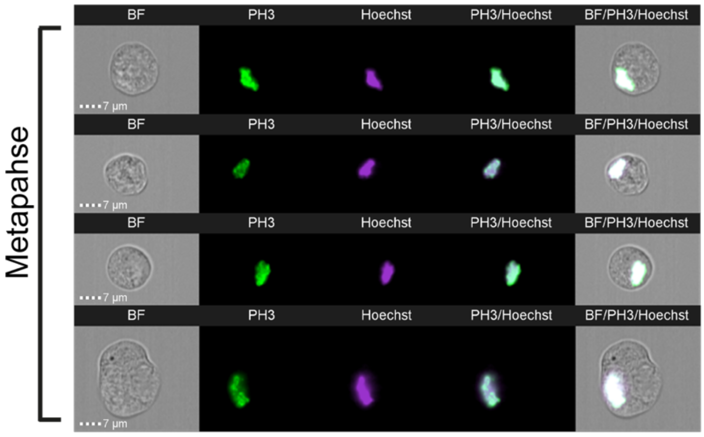


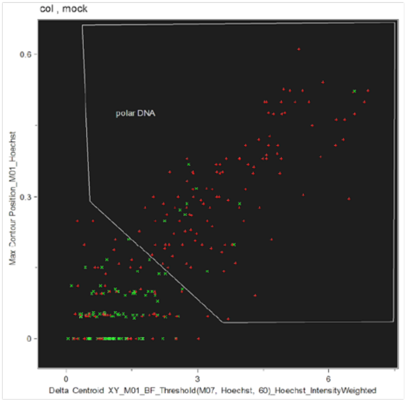


| Population | Count | %Gated |
|---|---|---|
| Colchicine treatment | 167 | 100 |
| Polar DNA | 107 | 64.1 |
| mock | 103 | 100 |
| Polar DNA | 10 | 9.71 |

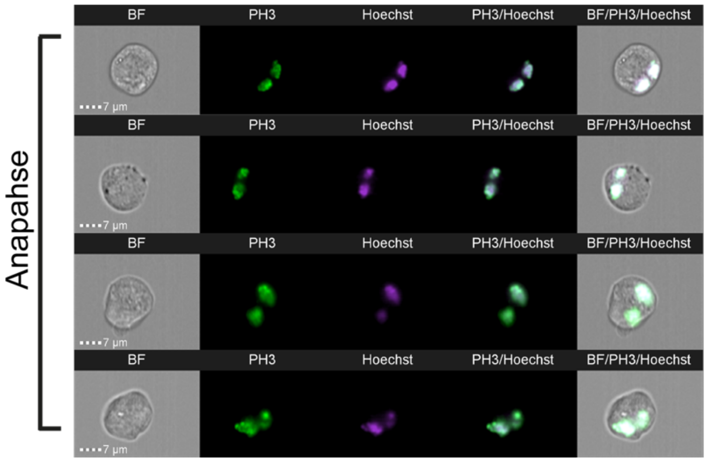


**Fig. S4.** *Drosophila* S2R+ cells treated with Colchicine display aberrant chromosomal morphology. Colchicine-treated cells were fixed with 4% PFA and stained with a phospho-Histone H3 (Ser10) antibody (green), Hoechst (DNA visualization; blue) and

Acti-stain 670 Phalloidin (filamentous Actin visualization; red). (a) Shown are representative images of cells in different mitotic phases obtained by imaging flow cytometry analysis. (b) Quantitation of cells with aberrant DNA morphology within mock and colchicine-treated cells using imaging flow cytometry analysis. Prophase- and metaphase-gated cells within either mock or colchicine-treated S2R+ cells, were further gated using calculations of the following features:
1. The Delta centroid XY feature (which measures the distance in microns between the centroid feature of two images using the user provided masks) was calculated using the BF default mask and Hoechst channel mask of the 60% most highly intense pixels. Cells with centered nucleus will get a lower value while polar located nucleus will get a higher value. 2. The Max contour position feature (the location of the contour in the cell that has the highest intensity concentration; the score is between 0 to 1, with 0 being the object center and 1 the object perimeter). To distinguish between central vs. polar location of the dividing nucleus, the Delta-centroid XY was plotted against the Max contour position, and polar DNA was gated as having the highest values of both features. Depicted are the analyses of cells in prophase and metaphase. Cells in anaphase are not shown due to the low number of cells observed in anaphase (Table S2).

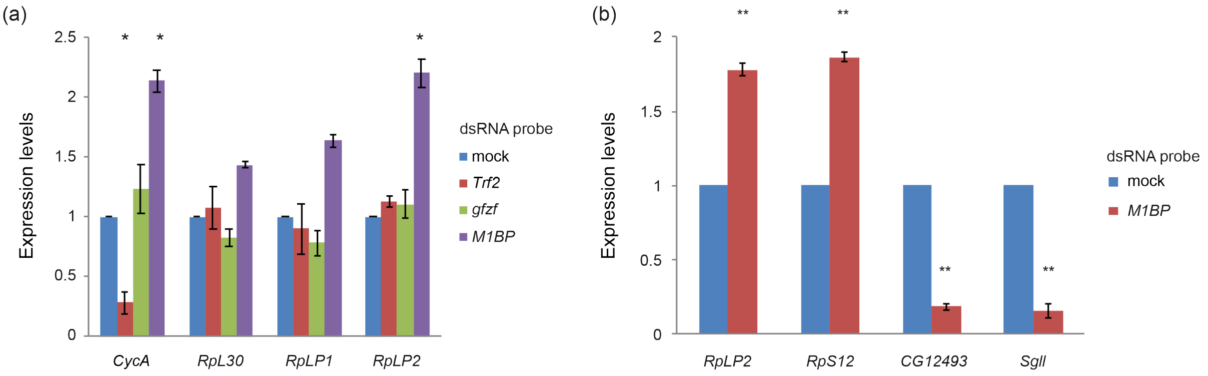


**Fig. S5.** Knockdown of *M1BP* specifically elevates the expression of *CycA* and ribosomal protein genes. *Drosophila* S2R+ cells were incubated for three days with dsRNA probes as indicated in the legend. RNA was isolated from the cells and reverse transcribed (RT) to cDNA. Real-time PCR (qPCR) experiments were used to analyze the RNA levels of the endogenous genes. qPCR experiments were performed in triplicates, and the graph represents the average of 3 experiments ± SEM. (a) *CycA* expression levels are reduced following *Trf2* knockdown, but not *gfzf* or *M1BP* knockdown. *p< 0.05, one-way ANOVA followed by Tukey's post hoc test as compared to the mock treatment of the relevant gene. (b) Additional ribosomal genes are influenced by *M1BP* knockdown, however not all genes are upregulated in response to *M1BP* knockdown. *CG12493* and *sgll* genes were previously shown to be downregulated upon *M1BP* knockdown [3].**p< 0.01, two-tailed Students t-test; compared to mock treatment.

**Table S1 (separate Excel file)**
GO terms analysis of the RNA-seq data. Summary, as well as the exact GO terms are provided, according to the datasheet name. Analysis was performed sing STRING. Only genes with pAdj <0.1 were considered. DEseq2 output is presented for either *Trf2* or *Tbp* knockdown, as compared to mock at 72h post silencing.

**Table S3 (separate Excel file)**
Mass-spectrometry data for short TRF2, long TRF2 and TBP proteins.

**Table S2.** Number of cells in each mitotic phase.
(a) Number of cells in each mitotic phase within phospho-Histone H3 Ser10-positive mitotic cells.

| | **Experiment #1** | | | **Experiment #2** | | | **Experiment #3** | | |
|---|---|---|---|---|---|---|---|---|---|
| **mitotic phase** | **P** | **M** | **A** | **P** | **M** | **A** | **P** | **M** | **A** |
| mock | 361 | 183 | 13 | 202 | 157 | 21 | 315 | 143 | 20 |
| Trf2 Probe #1 | 63 | 51 | 8 | 84 | 51 | 18 | 139 | 68 | 19 |
| *Trf2* Probe #2 | 80 | 38 | 8 | 86 | 51 | 10 | 93 | 56 | 20 |
| *Tbp* probe | 155 | 68 | 5 | 171 | 110 | 16 | 234 | 140 | 7 |
| Colchicine | 229 | 115 | 41 | 371 | 166 | 54 | 977 | 443 | 103 |

P=prophase, M=metaphase, A=anaphase.

(b) Number of cells in telophase. Phospho-Histone H3 Ser10-negative cells undergoing mitosis (as identified by the imaging flow cytometer) were defined as cells in telophase. It is of note that these cell counts are likely to include phospho-Histone H3 Ser10-negative doublet cells, which, even using the high-resolution Imagestream, could not be distinguished from telophase cells.

| | **Experiment #1** | **Experiment #2** | **Experiment #3** |
|---|---|---|---|
| **mitotic phase** | **T** | **T** | **T** |
| mock | 675 | 1489 | 837 |
| Trf2 Probe #1 | 361 | 1478 | 918 |
| *Trf2* Probe #2 | 575 | 1382 | 941 |
| *Tbp* probe | 299 | 1330 | 1029 |
| Colchicine | 288 | 1041 | 514 |

T=telophase
